# Developmental Arrest Associated with Altered Cerebellar Metabolism in Sudden Infant Death Syndrome

**DOI:** 10.64898/2026.07.20.739601

**Authors:** Javid Ghaemmaghami, Gabriele Simonti, Georgios Sanidas, Nora Wolff, Maria Triantafyllou, Rodolfo Cardenas Trejo, Henrik Sahlin Pettersen, Vittorio Gallo, Ioannis Koutroulis, Panagiotis Kratimenos

## Abstract

**Background:** Effective autoresuscitation during hypoxic stress demands the cerebellum integrate autonomic and arousal responses. It remains unknown if the SIDS cerebellum possesses the metabolic stability and developmental maturity to sustain this function. We aimed to define if cerebellar dysfunction constitutes a silent failure point within the central respiratory network.

**Methods:** We performed multi-omic integration of RNA-sequencing, targeted metabolomics, and quantitative histology on postmortem cerebellar tissue from SIDS infants and age-matched controls. Cross-omic concordance analysis mapped the relationship between metabolite abundance and the transcriptional directionality of corresponding metabolic enzymes.

**Results:** The SIDS cerebellum revealed a distinct profile defined by neuroinflammation and altered GABA and glutamate metabolism. Targeted metabolomics confirmed a metabolic imbalance with significantly elevated glutamate and GABA (*P* < 0.05). Quantitative histology exposed concurrent developmental arrest; persistent external granule layers highlighted morphological immaturity, explaining the failure to mount effective compensatory responses.

**Conclusions:** Metabolic pathology in the SIDS cerebellum is intrinsically linked to structural immaturity. This developmental arrest creates a compromised environment lacking the metabolic competence to support neural homeostasis, preventing critical arousal responses required for survival.

**Impact:**

- Multi-omic and neuropathological analysis reveal that structural immaturity in SIDS cerebellum is directly linked to neuroinflammation and metabolic excitotoxicity.
- Developmental arrest leaves the tissue metabolically incompetent to support the neural homeostasis required for infant survival during hypoxic stress.
- This study shifts the focus of SIDS pathology beyond traditional brainstem mechanisms by establishing the cerebellum as a critical, point of failure within the central respiratory network, providing a direct link between morphological delay and functional neurochemical disruption
- These findings offer translatable paradigm for understanding SIDS vulnerability and identify specific metabolic targets for future diagnostic risk screening and therapeutic interventions.

## INTRODUCTION

Deficits in the neural circuits regulating cardiorespiratory homeostasis and sleep-wake arousal represent a central neuropathological mechanism in sudden infant death syndrome (SIDS)^1–3^. Although the brainstem has long been implicated in SIDS, recent studies indicate that the cerebellar cortex also plays a critical role in respiratory recovery and autonomic defense^1,2,4,5^. This is consistent with the peak incidence of SIDS at 2–4 months of age, coinciding with a period of heightened vulnerability during cerebellar maturation, when granule cell precursors migrate from the external to the internal granule layer^6–9^. This migration is required for establishing the excitatory-inhibitory neurotransmitter balance essential for survival ^9–11^.

This critical phase of brain development is metabolically demanding^12^. An analysis of SIDS tissues shows a signature of inflammation and hypoxic stress; however, whether this dysregulation physically alters circuit architecture is unknown^13–15^. We hypothesize that the inflammatory and hypoxic milieu is not a transient response but rather stalls progenitor migration, resulting in structural developmental arrest and an anatomically immature cerebellar cortex.

Here, we correlated cellular and metabolic changes with cerebellar cytoarchitecture. We quantified cerebellar external granule layer retention and internal granule layer depletion to assess the developmental delay and analyzed Purkinje cell morphology to assess maladaptive plasticity. Linking these structural anomalies to established glutamatergic and GABAergic imbalances revealed a mechanism whereby developmental arrest of cerebellar circuits contributes to compensatory failure by preventing the establishment of the inhibitory tone required for recovery from transient respiratory challenges during sleep^4,5,16^.

## METHODS

### Human postmortem cohort tissue

This study was conducted in accordance with protocols approved by the Institutional Review Board of Children’s National Research Institute (IRB protocol #00015350). All postmortem subjects were from the Children’s National Hospital pathology department and the NIH Neurobiobank. Subjects were selected based on diagnostic classification; gestational age at birth and postmenstrual age at death; tissue availability; and preservation quality. All subject information is presented in Supplementary Table S3. Subjects with brain-destructive pathology (high-grade intraventricular hemorrhage, post-hemorrhagic hydrocephalus, cerebellar hemorrhage, or cystic periventricular leukomalacia) were excluded. Clinical variables, including prematurity, postnatal age, and relevant comorbidities, were recorded for downstream analysis. After legal pronouncement of death, an autopsy was typically initiated within 24-48 hours to limit postmortem degradation, with the postmortem interval reported for all tissue collected within this timeframe. Fixed-frozen cerebellar tissues were obtained at autopsy from the Department of Pathology at Children’s National Hospital. For formalin-fixed, paraffin-embedded (FFPE) samples, the brain was removed *en bloc*, weighed, inspected, and immediately immersed in 10% neutral-buffered formalin for at least 7 days to ensure uniform fixation. After fixation, the cerebellum was separated sharply from the brainstem at the level of the superior cerebellar peduncles and bisected precisely along the midline. Tissue sections were processed as FFPE specimens in accordance with College of American Pathologists-accredited histopathology protocols. Fixed tissue underwent graded ethanol dehydration, xylene clearing, and paraffin infiltration under vacuum before being embedded in fresh paraffin with standardized orientation to maintain a perpendicular cutting plane across the folial axis. Blocks were stored in a temperature- and humidity-controlled archive until microtomy. Sections were cut at 5 µm thickness on a rotary microtome using ribbons generated orthogonally to the folial curvature to ensure consistent representation of the external granule layer, molecular layer, internal granule layer, Purkinje cell layer, and deep white matter. Sections were floated on a 40°C water bath, mounted on positively charged slides, dried, and stained with hematoxylin and eosin (H&E).

### Mass spectrometry metabolomics

Targeted metabolomics was performed on flash-frozen postmortem human cerebellar tissue from 22 SIDS and 16 non-SIDS samples. Tissue samples were frozen before processing to preserve metabolic integrity. Neurotransmitters and pathway-associated analytes, including metabolites of the serotonin and dopamine pathways, norepinephrine, epinephrine, glutamate, GABA, and kynurenine, were analyzed by targeted mass spectrometry at the Vanderbilt Neurochemistry Core. Metabolite abundance was quantified from mass spectrometry signal intensities using ^13^C-labeled standards and normalized to tissue protein content. Final metabolite concentrations (ng/mg protein) were used for downstream statistical analysis.

### Bulk RNA-seq, pathway enrichment, and metabolomics analysis

Bulk RNA-seq was performed on flash-frozen postmortem human cerebellar tissue from 11 SIDS cases and 7 non-SIDS controls. Tissue was frozen before processing to preserve RNA integrity. Total RNA was extracted from cerebellar tissue using a standard RNA isolation workflow, followed by RNA quality assessment and library preparation for bulk transcriptomic profiling. Sequencing libraries were generated from extracted RNA and sequenced using a high-throughput short-read sequencing platform. Raw sequencing data were quantified using RSEM to generate transcript-level abundance estimates. Transcripts were summarized as gene-level counts and used for downstream differential expression analysis. Gene-level count distributions were examined across samples as an initial quality-control assessment and were broadly comparable between SIDS and non-SIDS samples (Fig. S1). Differential gene expression was performed using DESeq2. Gene counts were normalized using DESeq2 size-factor normalization, and differential expression was modeled using negative binomial generalized linear models. The primary model compared SIDS and non-SIDS cerebellar samples using SIDS status as the main variable. Log2 fold-change estimates were shrinkage-adjusted using the ashr method. Differentially expressed genes were defined using an FDR-adjusted p-value of < 0.05 and an absolute log2 fold change of > 0.585, corresponding to a 1.5-fold change. Because cerebellar gene expression is strongly influenced by developmental timing, a postmenstrual age-adjusted differential expression model was also fitted using the design formula ∼ postmenstrual age + SIDS status. This model tested the effect of SIDS status while including postmenstrual age at death as a continuous covariate, thereby accounting for transcriptional variation associated with developmental age. Principal component analysis was performed on variance stabilized expression values. Postmenstrual age was not regressed out of the expression matrix for this PCA; instead, samples were colored according to postmenstrual age. Differential-expression heatmaps were generated using genes identified from the postmenstrual age-adjusted model. Thus, gene selection was based on the SIDS effect estimated after accounting for postmenstrual age, whereas the heatmap displayed normalized, variance-stabilized expression values for those selected genes rather than expression values with postmenstrual age mathematically removed. Heatmaps were produced for genes meeting an FDR-adjusted p-value threshold of < 0.05 and, separately, for genes meeting a raw p-value threshold of < 0.05, both with and without dendrograms. Differential expression results were used for downstream pathway enrichment analyses. Pathway enrichment was performed using both over-representation analysis and gene set enrichment analysis. Over-representation analysis was conducted on significant differentially expressed genes using Hallmark, Gene Ontology Biological Process, KEGG, and Reactome pathway collections. Gene set enrichment analysis was performed using ranked gene-level statistics from the postmenstrual age-adjusted differential expression model across the same pathway databases. The pathway analysis used significance thresholds of FDR-adjusted p-value < 0.05 and absolute log2 fold change > 0.585 for defining significant differentially expressed genes. Transcriptomic and metabolomic findings were integrated at the pathway level because the RNA-seq and metabolomics datasets were generated from separate cohorts. Metabolite-associated pathways were compared with RNA-seq gene set enrichment results to identify biological themes showing concordant alteration across modalities. In addition, enzyme-level convergence analysis was performed for GABA and glutamate by mapping each metabolite to the genes encoding enzymes relevant to synthesis, degradation, and reuptake. SIDS versus non-SIDS RNA-seq log_2_ fold changes and p-values were extracted for the genes for these enzymes and visualized by enzyme role. This enzyme-level analysis was interpreted as exploratory because the enzyme genes did not reach FDR significance but provided directional context for the metabolite findings.

### Neuropathological analysis

Postmortem FFPE human cerebellar tissue from 22 SIDS and 24 non-SIDS infants was used for neuropathological analysis, matched by postmenstrual age. Purkinje cell density was quantified from cerebellar sections using a deep learning–based histopathology pipeline. Whole-slide imaging of H&E-stained sections was performed using an Olympus VS120 microscope at 20× magnification. The entire tissue area on each slide was quantified using deep learning–based segmentation. Briefly, an active-learning NoCodeSeg/DeepLabV3+ workflow was used, with a scripted Python/PyTorch ensemble of three DeepLabV3+ models with ResNet-18 backbones. Ensemble predictions were reviewed and corrected by an expert neuropathologist in QuPath and Microscopy Image Browser, and corrected annotations were reincorporated through iterative training cycles until performance stabilized. Final Purkinje cell predictions were imported into QuPath and restricted to the Purkinje cell layer at the molecular layer–internal granule layer interface. Cells located more than 25 µm from this interface were excluded. Purkinje cell density was normalized to estimated Purkinje cell layer length, calculated from a 50 µm-wide band centered on the molecular layer boundary, extending 25 µm toward the molecular layer and 25 µm toward the internal granule layer. Because the band perimeter includes both inner and outer boundaries, it was divided by 2 to approximate the Purkinje cell layer centerline length. Purkinje cell density was reported as Purkinje cells per mm of estimated Purkinje cell layer length. For downstream analysis, Purkinje cell density was compared between SIDS and non-SIDS subjects and modeled as a function of postmenstrual age to account for developmental age and prematurity. This normalization was used to account for differences in folial perimeter and tissue sampling area across subjects. Purkinje cell dendritic morphology was assessed using calbindin-labeled cerebellar sections. In high-resolution images, five Purkinje cells were manually traced per subject using Neurolucida. Total dendritic arbor length was measured for each traced cell, and the average dendritic length per subject was used for downstream statistical analysis. Granule cell precursors and mature granule cells were quantified from representative cerebellar regions. For each subject, four representative regions of interest (ROIs) were selected, with the same ROI size across all samples, at 100x oil-immersion magnification (Fig. S3). Granule cell precursors were counted in the external granular layer, and mature granule cells were counted in the internal granular layer. Individual cells within each ROI were counted manually, and counts across the four ROIs were averaged to obtain a single mean granule cell precursor count and a single mean mature granule cell count per subject. Because postmortem infant cerebellar tissue is biologically heterogeneous and cell count measurements were not assumed to be normally distributed, group comparisons between SIDS and non-SIDS subjects were performed using non-parametric statistical tests. Kruskal–Wallis tests with Dunn’s post-hoc correction and Wilcoxon rank-sum tests were used where appropriate. Linear regression models incorporating postmenstrual age and group status were also used to determine whether SIDS-associated differences in cell density or morphology persisted after accounting for developmental age.

### Linear regression analysis of metabolomics and neuropathology

Linear regression models were used to test associations between SIDS status and metabolomic or histologic outcomes, accounting for developmental age. For targeted metabolomics, duplicate measurements from the same subject were averaged to obtain a single value per subject. Glutamate and GABA concentrations were modeled as outcomes using linear regression, with SIDS status or subgroup contrast as the primary predictor and gestational age plus postnatal age as covariates. Postmenstrual age was calculated as the infant’s gestational age at birth plus their current chronological (postnatal) age. Metabolite concentrations were normalized to tissue protein content and reported as ng/mg protein. Group comparisons were further assessed using t-tests and Kruskal–Wallis tests. For Purkinje cell analysis, postmenstrual age metadata were merged with the Purkinje cell count dataset, and subjects missing postmenstrual age were excluded. Group differences in Purkinje cell count were assessed using Kruskal–Wallis and Dunn’s tests. To evaluate age-associated Purkinje cell count trajectories, Purkinje cell count was modeled as a function of postmenstrual age, diagnostic group, and their interaction. Separate within-group linear models were also used to estimate age-associated Purkinje cell count slopes in SIDS and non-SIDS subjects. Results were summarized using regression coefficients, confidence intervals, p-values, and group-based visualizations.

## RESULTS

### Transcriptomics Reveal Prominent Neuroinflammatory Signaling in the SIDS Cerebellum

To determine the molecular processes at death in the SIDS cerebellum, we characterized the transcriptomes of fresh-frozen cerebellar tissue from SIDS and age-matched controls using bulk RNA sequencing (RNA-seq) (Supplementary Table S1). The SIDS cerebellum exhibited a transcriptional profile distinct from that of non-SIDS cerebella. By analyzing the differentially expressed genes (DEGs) using Gene Set Enrichment Analysis (GSEA), we identified 730 DEGs in human cerebellar tissue from SIDS cases compared to age-matched controls (adjusted P-value < 0.05) (**Fig. 1a**). Principal component analysis (PCA) of the transcriptome demonstrated clustering of SIDS cases with the primary variance driven by disease status, with developmental age as a covariate (see methods) (**Fig. 1b**). The SIDS signature is characterized by a broad inflammatory response, with upregulation of the genes encoding interferon-γ (normalized enrichment score [NES] > 2.2), interferon-α (NES > 2.0), and tumor necrosis factor alpha (TNFA) signaling via the NF-κB pathways (**Fig. 1c**). As immune activation typically requires reallocation of cellular resources, we determined whether this inflammatory state compromised basic developmental functions. Consistent with a resource-diverting stress response, there was downregulation of pathways critical for cellular proliferation and maintenance, including DNA repair and Myc targets (NES < -1.5). Thus, the SIDS cerebellum prioritizes immune defense at the expense of homeostatic growth and repair.

**Fig. 1:**
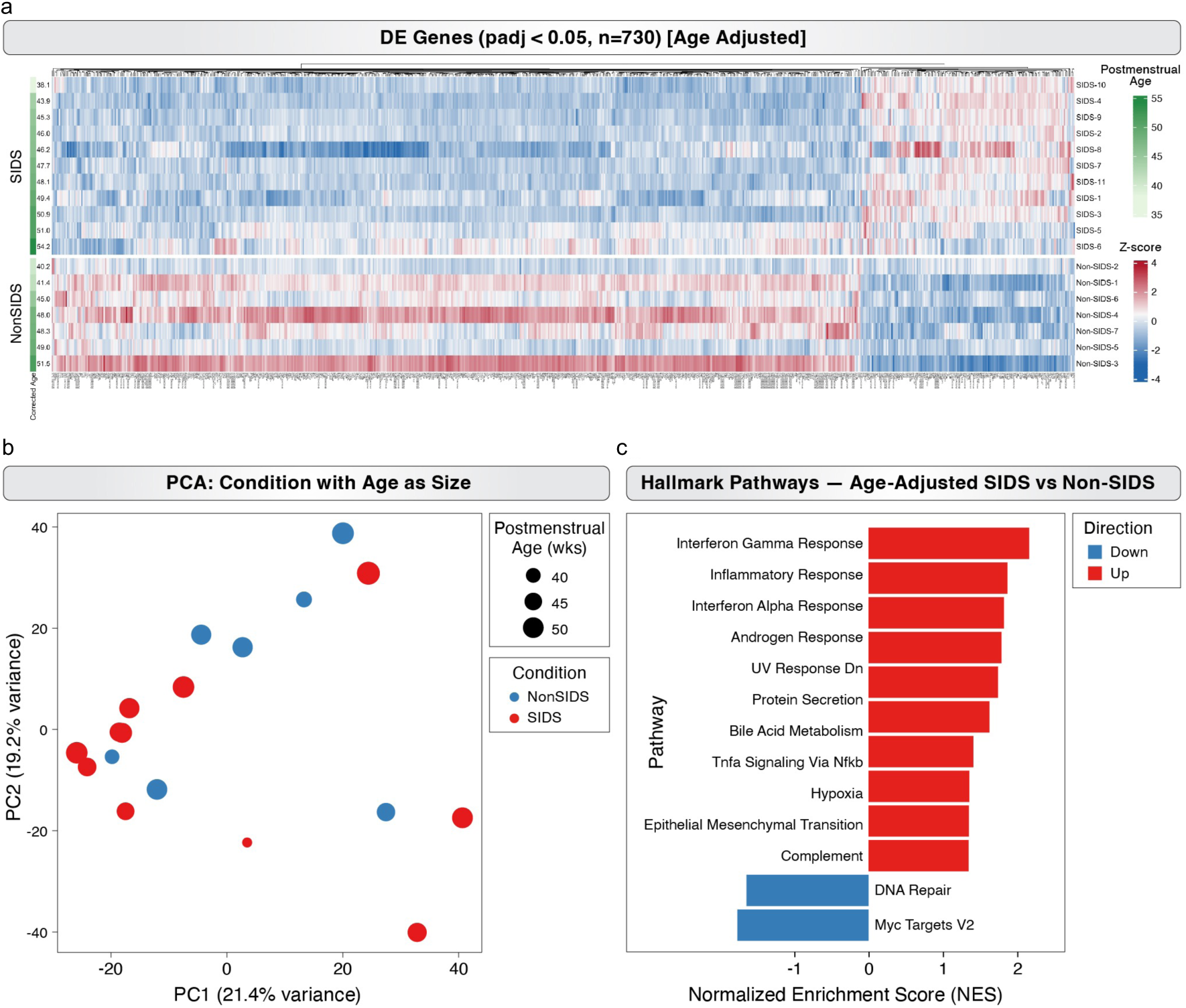
A global inflammatory transcriptional shift in the SIDS cerebellum. **a** Principal component analysis (PCA) of bulk RNA-seq transcriptomes from fresh-frozen human cerebellar tissue from SIDS (n = 11) and non-SIDS (n = 7) samples. Points are colored by diagnostic group and scaled by corrected age in weeks. PCA demonstrates separation between SIDS and non-SIDS cerebellar samples, with developmental age accounted for as a covariate in downstream analyses. **b** Gene set enrichment analysis (GSEA) of age-adjusted differentially expressed genes identifies enrichment of inflammatory and stress-associated pathways in the SIDS cerebellum, including interferon-γ response, inflammatory response, interferon-α response, TNFA signaling via NF-κB, hypoxia, and complement. Downregulated pathways include DNA repair and MYC targets. Bars indicate normalized enrichment score (NES) with red indicating upregulated pathways and blue indicating downregulated pathways in SIDS. **c** Heatmap of 730 differentially expressed genes identified between SIDS and non-SIDS cerebellar tissue using an age-adjusted differential expression model (padj < 0.05). Rows represent individual subjects grouped by diagnostic status, and columns represent differentially expressed genes. Z-scored expression values show transcriptional clustering between SIDS and non-SIDS samples after accounting for corrected age as a covariate.

### Targeted Metabolomics highlight Imbalance of Glutamate and GABA in SIDS Cerebellum

To determine whether the inflammatory environment in the SIDS cerebellum is associated with changes in neurochemical homeostasis, we quantified neurochemicals at the time of death using targeted metabolomics on fresh-frozen tissue (**Supplementary Table S2**, **Fig. S2**). To account for prematurity, we used postmenstrual age as an adjustment covariate in our linear models. postmenstrual age is calculated as an infant’s gestational age at birth plus their current chronological (postnatal) age, typically measured in weeks and days. We identified elevated levels of key neurotransmitters glutamate and GABA in SIDS cases compared to controls, with GABA levels increasing significantly with postmenstrual age (P = 0.039) (**Fig. 2a-b**). The elevated levels of both the primary excitatory and inhibitory transmitters indicate a widespread failure of synaptic clearance mechanisms. This accumulation indicates that the SIDS cerebellum operates in a persistently saturated metabolic state, which may compromise the precision of neural signaling required for respiratory control.

**Figure 2.**
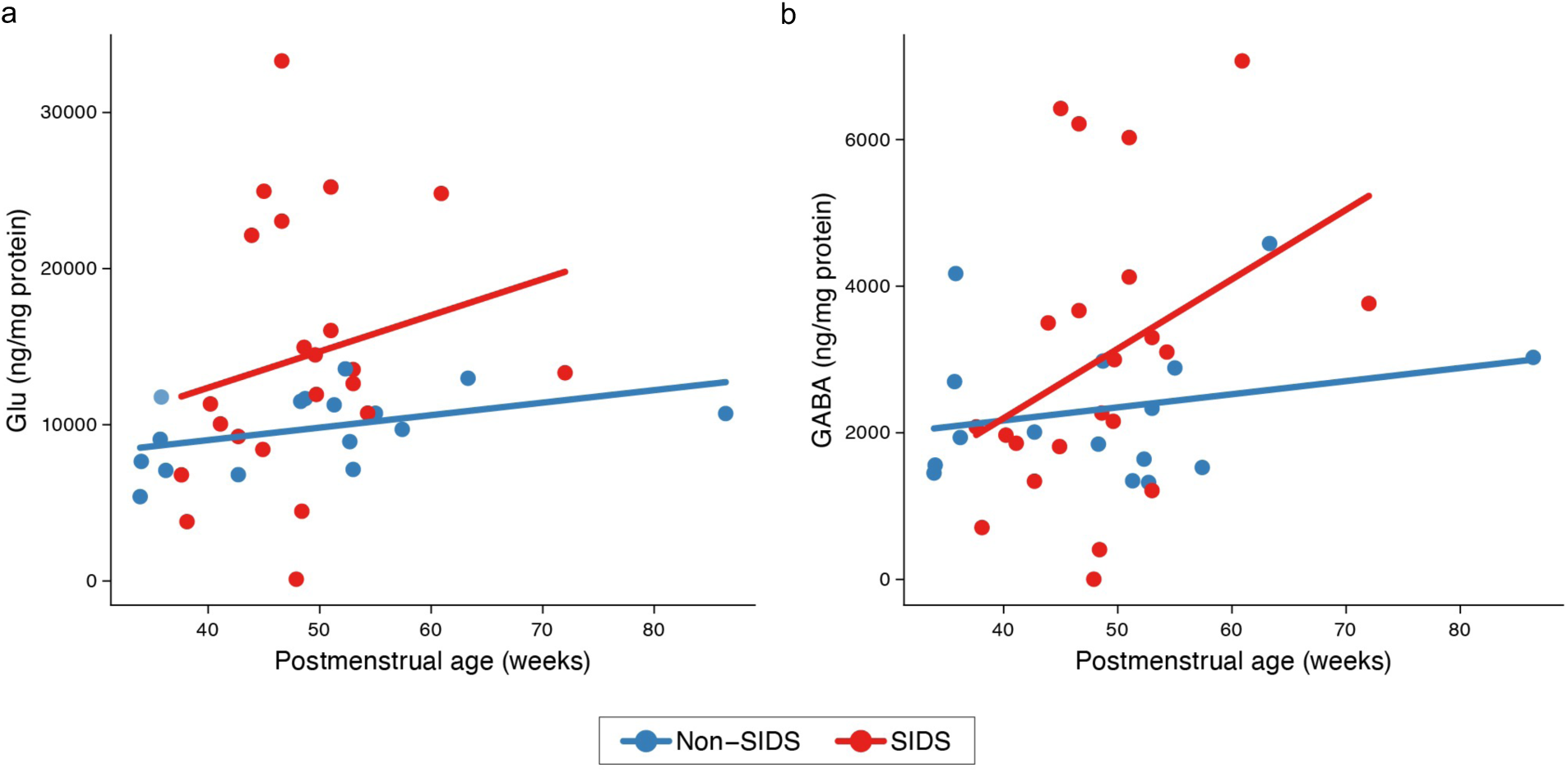
SIDS-associated age-related changes in glutamate and GABA in the cerebellum. **a** A scatter plot of cerebellar glutamate (ng/mg protein) vs. postmenstrual age (weeks) in non-SIDS controls (n = 17, blue) and SIDS cases (n = 22, red). Solid lines represent fitted linear regression trend lines showing the association between glutamate concentration and postmenstrual age within each group. Glutamate concentrations differed significantly between the SIDS and non-SIDS groups (P = 0.026). **b** A scatter plot of cerebellar GABA (ng/mg protein) as a function of postmenstrual age at the time of image (weeks) for non-SIDS controls (n = 17, blue) and SIDS cases (n = 22, red). Solid lines represent separate fitted linear regression trend lines showing the estimated association between GABA concentration and postmenstrual age within each group (P = 0.039).

### Cerebellar Metabolite Accumulation Aligns with Enzymatic Transcriptional Uncoupling

To determine whether the high glutamate and GABA levels reflected active upregulation of biosynthetic pathways or passive failure of clearance mechanisms, we compared metabolite abundance with the transcriptional activity of genes in their corresponding pathways. Concordance between RNA expression and metabolite levels would support transcriptional regulation, whereas discordance would suggest post-transcriptional or metabolic mechanisms such as clearance. Because the RNA-seq and metabolomics samples were from different subjects, we mapped each metabolite’s effect size (Cohen’s d) against the mean NES of matching RNA-seq GSEA pathways, thereby providing a significance-free assessment of cross-omic directional agreement.

We identified different responses in SIDS pathology (**Fig. 3a**). First, we identified a concordant relationship for the excitotoxins glutamate and GABA, as accumulation of both metabolites correlated with the upregulation of their associated signaling pathways. Thus, the high levels of glutamate and GABA in the SIDS cerebellum suggest that it is transcriptionally programmed to maintain this excitotoxic state. In contrast, serotonin accumulated despite reduced transcription of genes in the serotonin pathway. Thus, high serotonin levels are likely due to failure of clearance or degradation mechanisms rather than overproduction.

**Figure 3.**
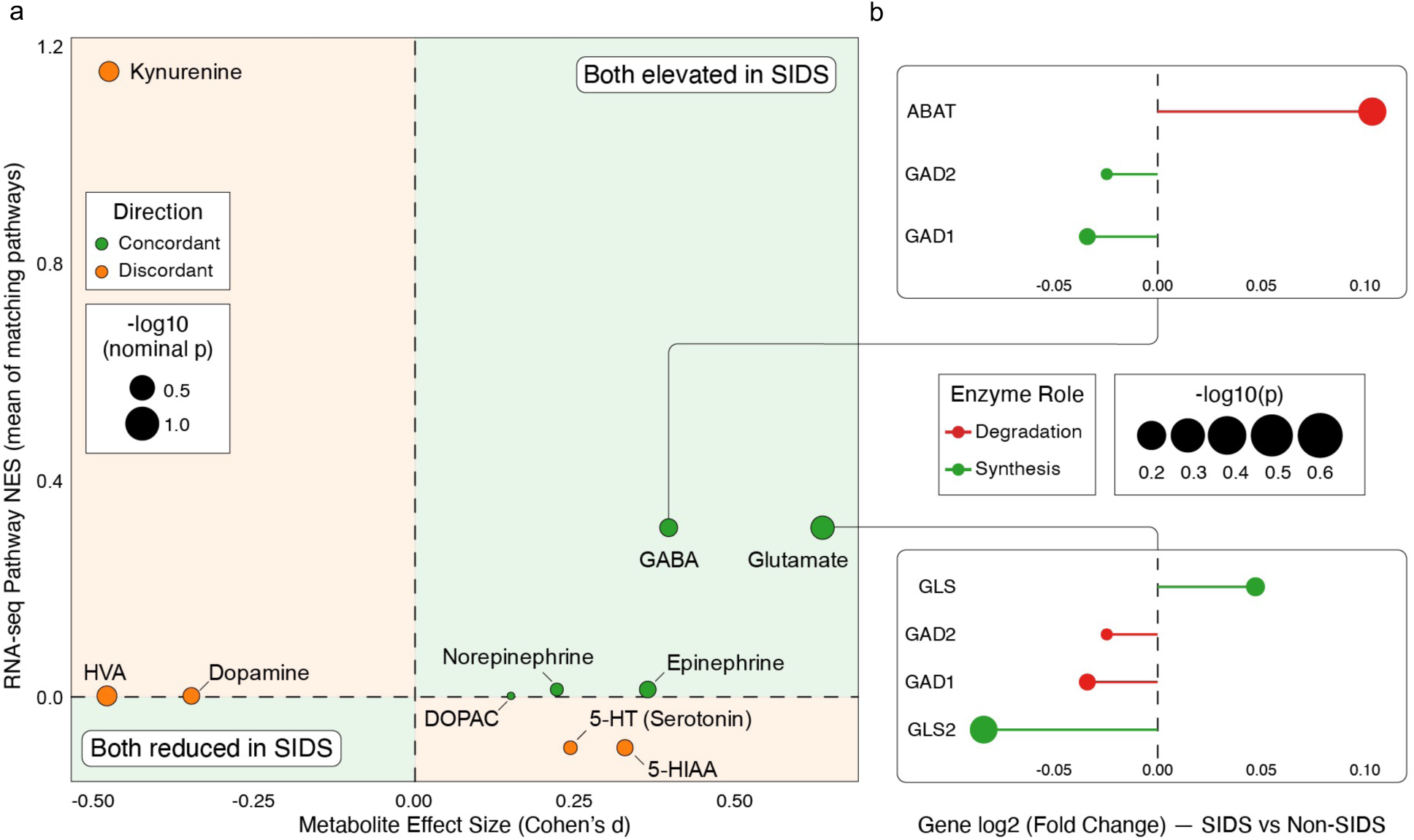
Dysregulated neurotransmitter pathways in SIDS. **a** A scatter plot integrating metabolite effect sizes (Cohen’s d, x-axis) of non-SIDS (n = 17) and SIDS (n = 22) cases with RNA-seq pathway normalized enrichment scores (NES, y-axis) derived from non-SIDS controls (n = 7) and SIDS cases (n = 11). Because metabolomics and RNA-seq were performed on separate subject cohorts, integration was performed at the pathway level rather than by subject-level matching. Green circles indicate concordant directionality between metabolomic and transcriptomic changes, whereas orange circles represent discordant pathways. Marker size reflects the statistical significance (-log_10_ nominal p-value). **b** Details of gene expression changes for key enzymes involved in GABA and glutamate metabolism in the same cohort. Horizontal dumbbell plots display the log_2_ fold change (SIDS vs. non-SIDS) for synthesis enzymes (green) and degradation enzymes (red), with marker size indicating statistical significance (-log_10_ p-value).

To identify the pathway genes that drive glutamate accumulation, we examined the regulation of its rate-limiting enzymes. The simultaneous upregulation of the gene encoding the glutamate-synthesizing enzyme GLS and the downregulation of the genes encoding the conversion enzymes GAD1 and GAD2 create a metabolic bottleneck (**Fig. 3b**). This enzymatic discordance confirms that the tissue is transcriptionally programmed to accumulate glutamate, thereby establishing a persistent excitotoxic environment that may result from the structural immaturity of the tissue.

### Cerebellar Cortical Morphology Reflects Metabolic and Molecular Changes in SIDS

To determine whether the inflammatory state and the increases in glutamatergic and GABAergic signaling were associated with cellular developmental arrest, we analyzed formalin-fixed, paraffin-embedded (FFPE) cerebellar tissues from 22 SIDS and 24 non-SIDS subjects, accounting for prematurity by assessing preterm subjects at term-equivalent postmenstrual age (**Supplementary Table S3**). Using the external granule layer persistence (as indicated by elevated GCP counts) and granule cell precursor retention as markers of maturation, we found a significant retention of granule cell precursors in the external granule layer (**Fig. 4a-b**) and depletion of differentiated granule cells in the internal granule layer (P = 0.024) (**Fig. 4c**), indicating a stalled granule cell migratory process. These changes were associated with a decreased Purkinje cell density in SIDS cases compared to controls (**Fig. 4d**). There was an age-associated decline in Purkinje cell count in both SIDS and non-SIDS subjects; however, the decline was steeper with postmenstrual age in non-SIDS subjects. To determine whether the surviving output neurons showed morphological adaptations to this environment, we quantified dendritic arborization by measuring total dendritic length, which was increased in SIDS Purkinje cells (P = 0.0018) (**Fig. 4e**). These reduced Purkinje cell numbers, coupled with dendritic elongation, suggest a compensatory response in surviving Purkinje cells, aligning with the need to maintain GABA secretion and accommodate increased glutamate input within a developmentally arrested circuit.

**Figure 4.**
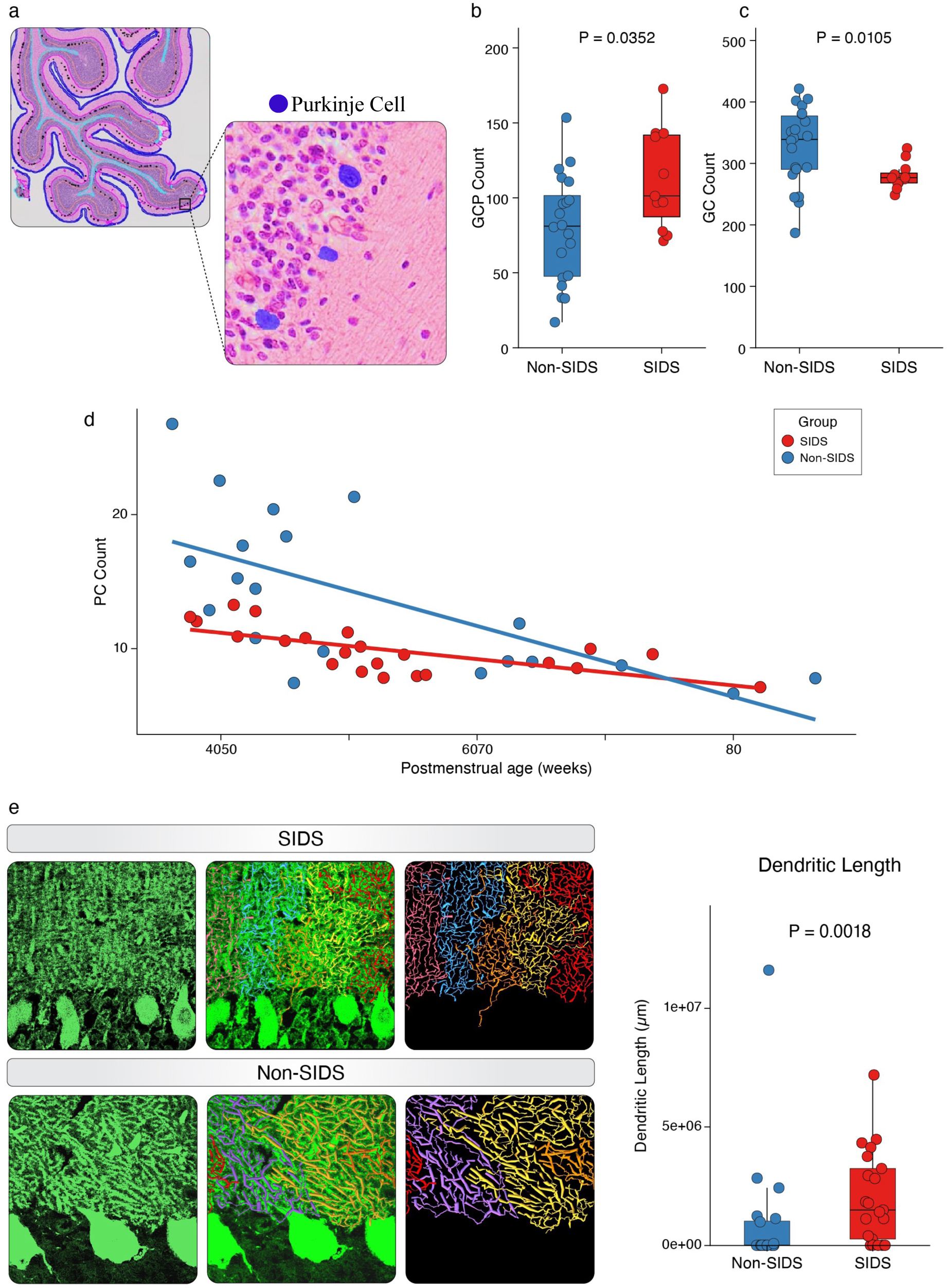
SIDS-associated changes in cerebellar cellular composition and dendritic morphology. **a** A representative image demonstrating the cortical segmentation used for quantification of Purkinje cells (PC), mature granule cells (GC), and granule cell precursors (GCP) in the cerebellum, generated by a whole-slide imaging-based automated algorithm. **b, c** Box plots of GCP counts (panel b) and GC counts (panel c) for non-SIDS controls (n = 23) and SIDS cases (n = 12). **d** A scatter plot of PC count vs. postmenstrual age (weeks) for non-SIDS controls (n = 24, blue) and SIDS cases (n = 22, red). Solid lines represent fitted linear regression trend lines showing the estimated association between PC count and postmenstrual age for each cohort. **e** Representative Neurolucida 360 images and the corresponding box plot comparing total dendritic length (micrometers) between SIDS cases (n = 22) and non-SIDS (n = 24) controls. All p-values are shown in the graphs.

## DISCUSSION

We describe here an inflammatory signature in the SIDS cerebellum and demonstrate its association with the structural developmental arrest of the cortical circuit. Although SIDS has historically been associated with medullary serotonergic dysregulation^1,3,17^, our data suggest that during the critical 2-4-month SIDS risk window^18^, the cerebellum may also have a role in this pathology. External granule layer retention in SIDS cerebellar tissue is similar to the histological phenotypes reported in models of chronic perinatal hypoxia^9,15,19^, suggesting that an inflammatory milieu drives the developmental arrest of granule cell migration. This arrest leaves the cerebellar cortex in an anatomically immature state, and potentially decoupled from the developing brainstem networks required for cardiorespiratory recovery^4,5,20^.

This arrest of granule cell migration was accompanied by structural remodeling of the cerebellar output circuitry, and the reduction in Purkinje cell density is consistent with the well-documented vulnerability of these neurons to inflammatory and hypoxic insults^21–23^. However, the surviving neurons exhibited marked dendritic hypertrophy, suggesting that they undergo structural modifications rather than simple neurodegeneration. We propose that this dendritic expansion is a homeostatic compensatory response to disrupted circuit development. Without robust granule cell input and under excitotoxic stress, surviving Purkinje cells integrate a broader range of synaptic inputs and partially preserve inhibitory output^24–26^. This interpretation provides a structural basis for the glutamatergic/GABAergic imbalance observed in our cohort, suggesting that the excitotoxicity is not merely chemical but a product of mismatched synaptic scaling.

This structural disintegration also appears to disrupt the wider monoaminergic network. We observed a significant accumulation of serotonin and 3,4-dihydroxyphenylacetic acid (DOPAC) in the cerebellar cortex. In contrast with reports of serotonergic depletion in the medullary raphe of SIDS victims^1,3,17,27^, our findings suggest a failure of the ponto-medullary-cerebellar loop. Impaired neurotransmitter release or altered axonal transport may lead to accumulation of neurotransmitters or their metabolites within cerebellar terminal fields^28,29^. Together, these findings suggest that the SIDS cerebellum is not only developmentally immature but also exhibits altered neurochemical communication with brainstem respiratory circuits.

These structural and neurochemical abnormalities may reduce the cerebellum’s ability to support respiratory and autonomic responses during hypoxic stress. The cerebellum modulates respiratory recovery via inhibitory projections to the fastigial nucleus^30–33^. The reduction in Purkinje cell density suggests that dendritic hypertrophy of the surviving neurons cannot fully preserve circuit function. Thus, SIDS may not represent an instantaneous failure of arousal but rather the collapse of a compensatory reserve. We propose that developmental arrest of cerebellar circuitry, together with inflammatory and neurochemical dysregulation, diminishes the contribution of the cerebellum to respiratory recovery during transient hypoxic events, thereby transforming a recoverable sleep apnea into terminal respiratory failure.

Our results expand the understood neuropathology of SIDS, suggesting that cerebellar dysregulation may act in conjunction with established brainstem abnormalities to compromise infant survival. We show that changes in the SIDS cerebellum correlate with a structural developmental delay, manifested in the retention of the external granule layer and a compensatory hypertrophy of surviving Purkinje cells. This structural vulnerability and the anomalous accumulation of serotonin suggest that the altered cerebellar circuit cannot supply the inhibitory modulation necessary to recover from sleep-related apnea. Thus, rather than acting as an isolated trigger, these cerebellar deficits likely integrate with brainstem dysfunction, culminating in a collective failure of protective autoresuscitation during the most vulnerable window of postnatal development.

### Limitations

Our findings were constrained by the inherent nature of postmortem neuropathological investigation. First, the sample size was restricted by the stringent inclusion criteria required for high-quality tissue preservation. This ensured the validity of our molecular assays but limited our statistical power to detect small differences between potential SIDS subgroups. Second, although we excluded cases with overt autolysis, the variability in the timing of postmortem tissue acquisition may have affected metabolite quantification, particularly for labile monoamines such as serotonin. Third, this cross-sectional study found a robust association between the inflammatory signature and structural arrest but did not establish causality. Although we identified inflammation within the critical risk window, we could not determine whether this immune activation preceded the developmental arrest. Finally, because clinical data on the subjects’ sleep environments were unavailable in all cases, we could not determine how exogenous stressors might interact with the molecular pathology described here.

## ACKNOWLEDGMENTS

We thank the CNH and NIH NeuroBioBank for providing postmortem cerebellar tissue and the families who consented to autopsy. We honor the memory of the deceased infants whose profound contribution has advanced our understanding of SIDS; through their gift, we strive to improve the quality of life for future generations of children.

## Funding

This work was supported by the Raynor Cerebellum Project (P.K.), American SIDS Institute (P.K.), a Children’s National Board of Visitors’ Grant (P.K.),

## Author contributions

J.G. performed experiments, analyzed and interpreted data, performed bioinformatic analyses and computational modeling, and wrote the manuscript. G.Si. and G.S. performed experimental work and contributed to data acquisition. N.W., M.T., and R.C.T. contributed to data acquisition. H.S.P. developed and implemented the deep learning–based neuropathological segmentation pipeline. I.K. contributed to data acquisition and manuscript revisions. V.G. contributed to manuscript revision. P.K. conceived and supervised the study, designed experiments, interpreted results, and wrote and edited the manuscript.

---

**Supplementary Figure 1.**
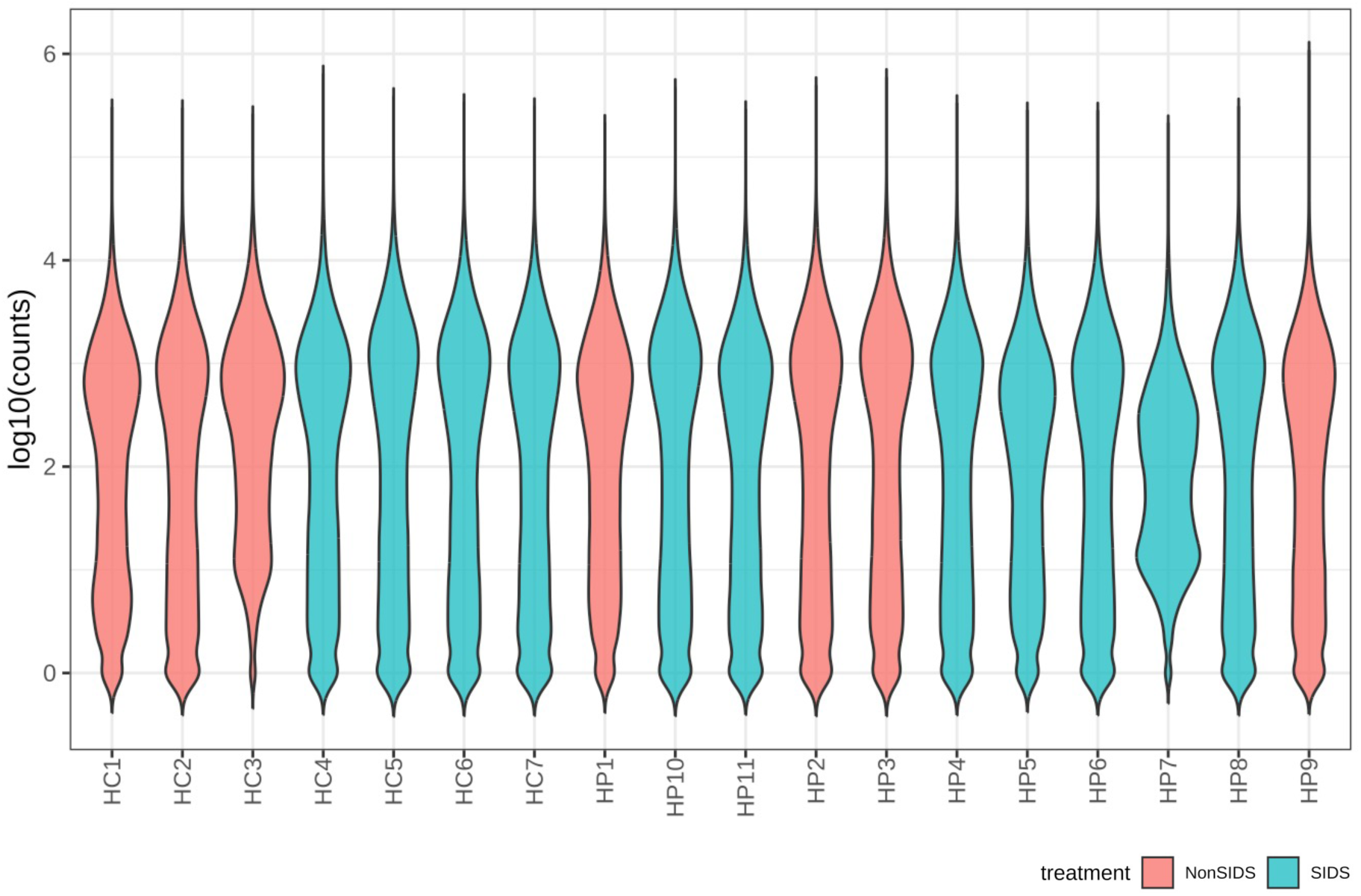
Distribution of gene level read counts across bulk RNA-sequencing samples. Violin plots show the distribution of log10-transformed gene counts for each cerebellar bulk RNA-seq sample. Samples are ordered along the x-axis and colored by treatment group: NonSIDS (salmon; *n* = 7) and SIDS (teal; *n* = 11). The width of each violin represents the relative density of genes at a given expression level. Similar count distributions across samples indicate broadly comparable sequencing depth and expression distribution profiles. This plot was used as an initial quality control assessment before downstream normalization and differential expression analysis.

**Supplementary Figure 2.**
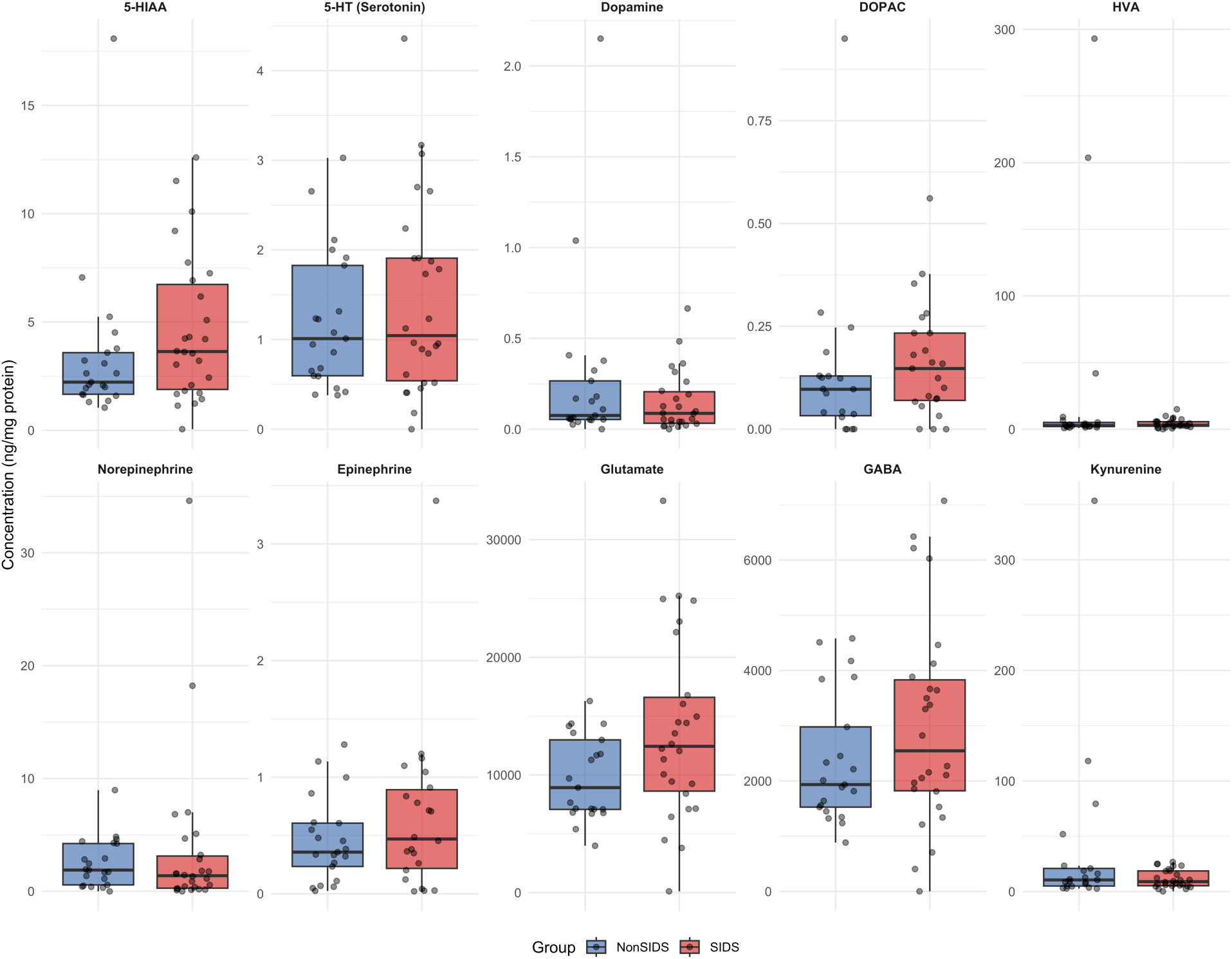
Neurotransmitter metabolite profiles in cerebellar tissue from SIDS and non-SIDS. Cerebellar concentrations of 10 neurotransmitter metabolites in non-SIDS and SIDS samples: 5-hydroxyindoleacetic acid (5-HIAA), serotonin [5-HT], dopamine, 3,4-dihydroxyphenylacetic acid (DOPAC), homovanillic acid (HVA), norepinephrine, epinephrine, glutamate, γ-aminobutyric acid (GABA), and kynurenine. Concentrations are expressed as ng per mg protein. Each point represents an individual sample; boxes indicate the interquartile range with the median shown by the central line, and whiskers represent the distribution beyond the quartiles. The metabolomics cohort comprised 16 non-SIDS and 22 SIDS samples.

**Supplementary Figure 3.**
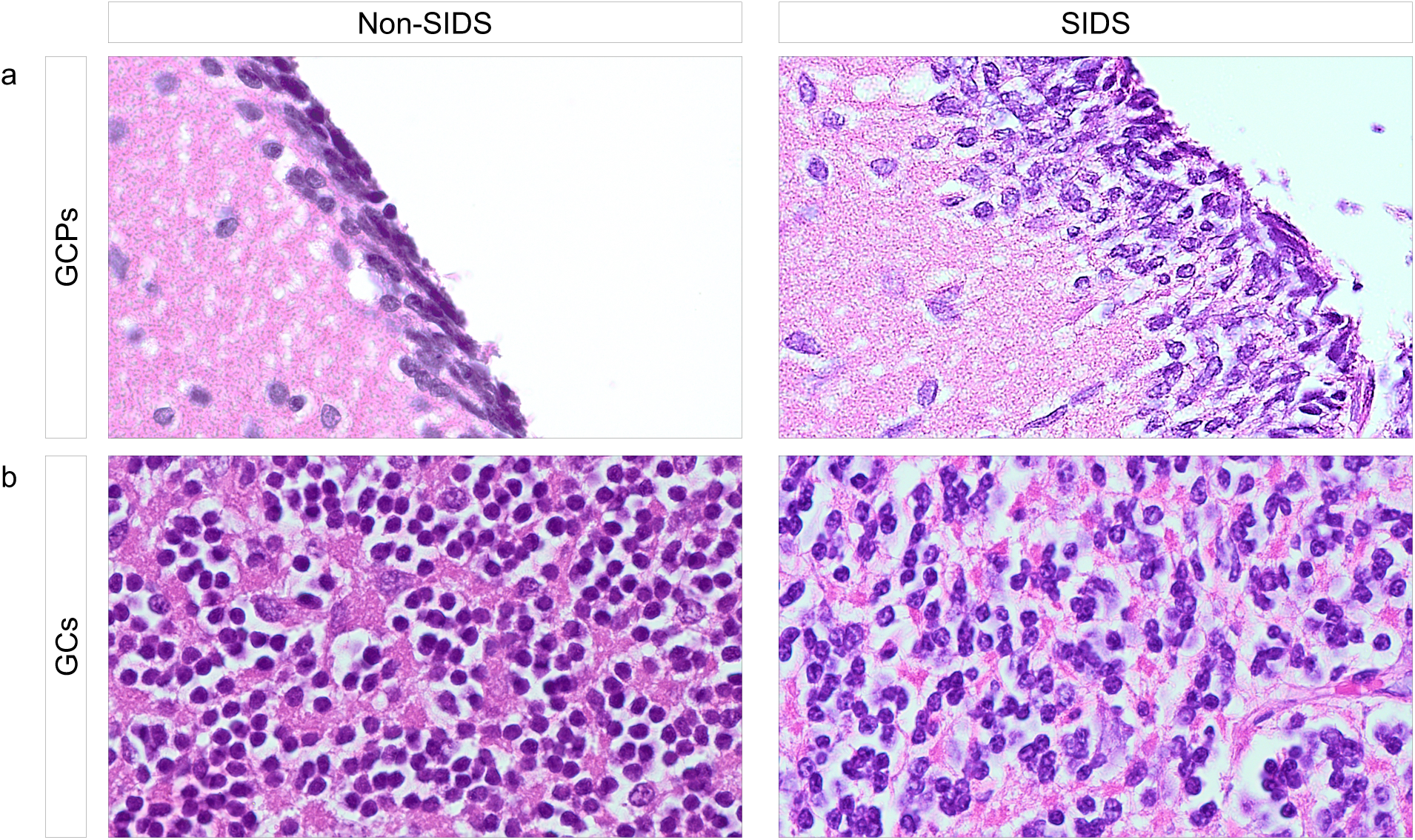
Representative 100x images of GCPs and GCs in SIDS and non-SIDS cerebellar tissue. **a,** Representative hematoxylin and eosin (H&E)-stained 100x images of GCPs from SIDS and non-SIDS samples. SIDS cases exhibited higher GCP counts than non-SIDS cases. **b,** Representative H&E-stained 100× images of GCs from SIDS and non-SIDS samples. GC counts were higher in non-SIDS cases than in SIDS cases.

**Supplementary Table S1.**
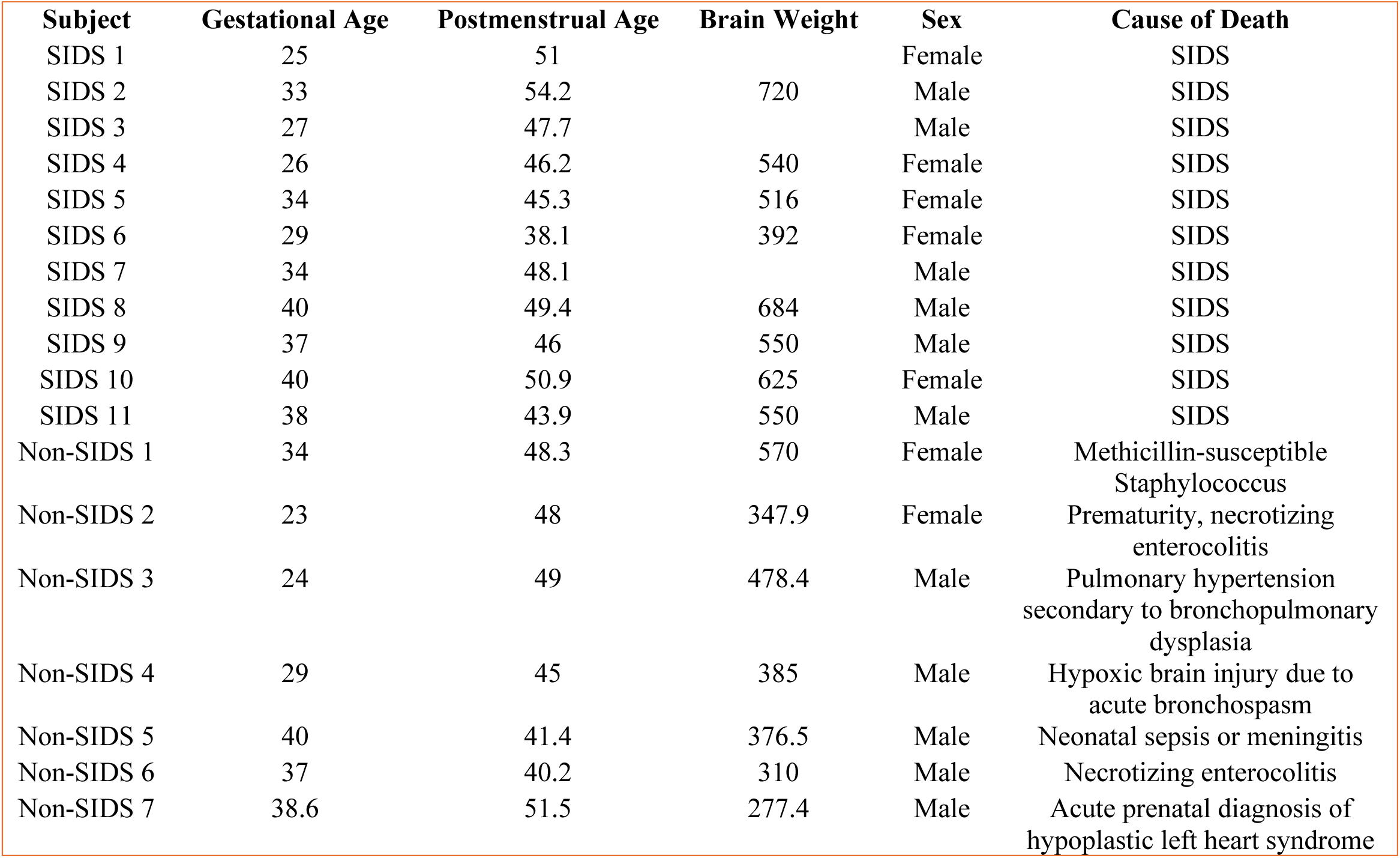
Subject demographics for bulk RNA-seq.

| **Subject** | **Gestational Age** | **Postmenstrual Age** | **Brain Weight** | **Sex** | **Cause of Death** |
| --- | --- | --- | --- | --- | --- |
| SIDS 1 | 25 | 51 |  | Female | SIDS |
| SIDS 2 | 33 | 54.2 | 720 | Male | SIDS |
| SIDS 3 | 27 | 47.7 |  | Male | SIDS |
| SIDS 4 | 26 | 46.2 | 540 | Female | SIDS |
| SIDS 5 | 34 | 45.3 | 516 | Female | SIDS |
| SIDS 6 | 29 | 38.1 | 392 | Female | SIDS |
| SIDS 7 | 34 | 48.1 |  | Male | SIDS |
| SIDS 8 | 40 | 49.4 | 684 | Male | SIDS |
| SIDS 9 | 37 | 46 | 550 | Male | SIDS |
| SIDS 10 | 40 | 50.9 | 625 | Female | SIDS |
| SIDS 11 | 38 | 43.9 | 550 | Male | SIDS |
| Non-SIDS 1 | 34 | 48.3 | 570 | Female | Methicillin-susceptible<br>Staphylococcus |
| Non-SIDS 2 | 23 | 48 | 347.9 | Female | Prematurity, necrotizing<br>enterocolitis |
| Non-SIDS 3 | 24 | 49 | 478.4 | Male | Pulmonary hypertension<br>secondary to bronchopulmonary<br>dysplasia |
| Non-SIDS 4 | 29 | 45 | 385 | Male | Hypoxic brain injury due to<br>acute bronchospasm |
| Non-SIDS 5 | 40 | 41.4 | 376.5 | Male | Neonatal sepsis or meningitis |
| Non-SIDS 6 | 37 | 40.2 | 310 | Male | Necrotizing enterocolitis |
| Non-SIDS 7 | 38.6 | 51.5 | 277.4 | Male | Acute prenatal diagnosis of<br>hypoplastic left heart syndrome |

**Supplementary Table S2.** Subject demographics for metabolomics analysis.

| **Subject** | **Gestational Age** | **Postmenstrual Age** | **Sex** | **Cause of Death** |
| --- | --- | --- | --- | --- |
| SIDS 1 | 33 | 37.6 | Male | SIDS |
| SIDS 2 | 28 | 38.1 | Male | SIDS |
| SIDS 3 | 40 | 40.2 | Male | SIDS |
| SIDS 4 | 35 | 41.1 | Female | SIDS |
| SIDS 5 | 29 | 42.7 | Male | SIDS |
| SIDS 6 | 40 | 43.9 | Female | SIDS |
| SIDS 7 | 40 | 44.9 | Male | SIDS |
| SIDS 8 | 32 | 45 | Male | SIDS |
| SIDS 9 | 40 | 46.6 | Male | SIDS |
| SIDS 10 | 26.3 | 46.6 | Female | SIDS |
| SIDS 11 | 39 | 47.9 | Male | SIDS |
| SIDS 12 | 39 | 48.4 | Male | SIDS |
| SIDS 13 | 33 | 48.6 | Female | SIDS |
| SIDS 14 | 38 | 49.6 | Male | SIDS |
| SIDS 15 | 35 | 49.7 | Male | SIDS |
| SIDS 16 | 39 | 51 | Female | SIDS |
| SIDS 17 | 25 | 51 | Female | SIDS |
| SIDS 18 | 40 | 53 | Female | SIDS |
| SIDS 19 | 35 | 53 | Female | SIDS |
| SIDS 20 | 33 | 54.3 | Male | SIDS |
| SIDS 21 | 32 | 60.9 | Female | SIDS |
| SIDS 22 | 40 | 72 | Male | SIDS |
| Non-SIDS 1 | 33 | 33.9 | Female | Acute bronchopneumonia |
| Non-SIDS 2 | 34 | 34 | Female | Bronchopneumonia |
| Non-SIDS 3 | 34 | 35.6 | Male | Anoxic encephalopathy due to drowning |
| Non-SIDS 4 | 35 | 35.8 | Female | Asphyxia |
| Non-SIDS 5 | 36 | 36.2 | Female | Complications of disorders |
| Non-SIDS 6 | 30 | 42.7 | Female | Viral syndrome with focal acute pneumonia |
| Non-SIDS 7 | 34 | 48.3 | Female | Prematurity associated |
| Non-SIDS 8 | 39 | 48.7 | Male | Methicillin-susceptible Staphylococcus |
| Non-SIDS 9 | 29 | 51.3 | Female | Multisystem failure |
| Non-SIDS 10 | 40 | 52.3 | Male | Bronchopneumonia |
| Non-SIDS 11 | 39 | 52.7 | Male | Hypoplastic left heart syndrome |
| Non-SIDS 12 | 38 | 53 | Male | Asphyxia |
| Non-SIDS 13 | 38 | 55 | Female | Reversed arterial perfusion syndrome (TRAP) |
| Non-SIDS 14 | 31 | 57.4 | Male | Hypoplastic left heart syndrome |
| Non-SIDS 15 | 30 | 63.3 | Male | Prematurity, acute respiratory failure |
| Non-SIDS 16 | 39 | 86.4 | Male | Prematurity associated w/ maternal eclampsia |

**Supplementary Table S3.** Subject demographics for neuropathological analysis and histology.

| **Subject** | **Gestational Age** | **Postmenstrual Age** | **Brain Weight** | **Sex** | **Cause of Death** |
| --- | --- | --- | --- | --- | --- |
| SIDS 1 | 25 | 51 |  | Female | SIDS |
| SIDS 2 | 27 | 46.4 |  | Male | SIDS |
| SIDS 3 | 28 | 38.1 | 392 | Male | SIDS |
| SIDS 4 | 29 | 42.7 | 470 | Male | SIDS |
| SIDS 5 | 32 | 60.9 | 661 | Female | SIDS |
| SIDS 6 | 32 | 45 | 535 | Male | SIDS |
| SIDS 7 | 33 | 37.6 | 460 | Male | SIDS |
| SIDS 8 | 33 | 57.4 | 720 | Male | SIDS |
| SIDS 9 | 35 | 55.3 | 671 | Female | SIDS |
| SIDS 10 | 35 | 41.1 | 512 | Female | SIDS |
| SIDS 11 | 35 | 49.7 | 650 | Male | SIDS |
| SIDS 12 | 39 | 48.7 | 576 | Male | SIDS |
| SIDS 13 | 39 | 52.2 | 600 | Female | SIDS |
| SIDS 14 | 39 | 82.1 | 804 | Male | SIDS |
| SIDS 15 | 39 | 56 | 720 | Female | SIDS |
| SIDS 16 | 40 | 41.3 | 380 | Female | SIDS |
| SIDS 17 | 40 | 68.85 |  | Male | SIDS |
| SIDS 18 | 40 | 73.7 | 783 | Female | SIDS |
| SIDS 19 | 40 | 50.9 | 539 | Male | SIDS |
| SIDS 20 | 40 | 52.7 |  | Male | SIDS |
| SIDS 21 | 40 | 65.6 | 930 | Male | SIDS |
| SIDS 22 | 41 | 49.9 |  | Male | SIDS |
| Non-SIDS 1 | 23 | 41.3 | 368 | Male | ARDS |
| Non-SIDS 2 | 23 | 55 | 275 | Female | Acute respiratory failure |
| Non-SIDS 3 | 24 | 64.3 |  | Female | Respiratory insufficiency |
| Non-SIDS 4 | 24 | 48 | 478.4 | Male | Pulmonary distension |
| Non-SIDS 5 | 25 | 37.6 | 310 | Male | NEC |
| Non-SIDS 6 | 29 | 45.1 | 385 | Male | Hypoxic injury |
| Non-SIDS 7 | 29 | 51.3 | 600 | Female | Viral syndrome/pneumonia |
| Non-SIDS 8 | 30 | 63.3 | 576 | Male | Pulmonary hemorrhage secondary to prematurity |
| Non-SIDS 9 | 30 | 42.7 | 380 | Female | Complications of prematurity |
| Non-SIDS 10 | 32 | 80.7 | 806 | Male | Epilepsy NOS |
| Non-SIDS 11 | 34 | 48.3 | 570 | Female | Methicillin-susceptible Staphylococcus |
| Non-SIDS 12 | 35 | 36.2 | 288 | Female | Hypoplastic left heart syndrome |
| Non-SIDS 13 | 38.7 | 39.1 | 400 | Male | Hypoplastic lungs |
| Non-SIDS 14 | 39 | 60.3 | 920 | Female | Congenital biliary atresia |
| Non-SIDS 15 | 39 | 42.4 |  | Female | Acute respiratory failure |
| Non-SIDS 16 | 39 | 86.4 | 1050 | Male | Drowned |
| Non-SIDS 17 | 39 | 44.1 | 339 | Female | Congenital heart disease |
| Non-SIDS 18 | 39 | 71.28 | 710 | Male | Acute multiorgan system failure |
| Non-SIDS 19 | 39 | 41.7 |  | Female | Viral myocarditis |
| Non-SIDS 20 | 39 | 45.7 | 1595 | Male | Multiorgan System Failure |
| Non-SIDS 21 | 39 | 62.4 |  | Female | Pneumonia |
| Non-SIDS 22 | 39 | 62.7 | 1140 | Male | Complication of the disorder |
| Non-SIDS 23 | 39.3 | 39.9 | 340 | Male | Infarct |
| Non-SIDS 24 | 40 | 42.7 | 224 | Male | Bronchopneumonia |

## Notes

### Competing Interest Statement

The authors have declared no competing interest.

